# Rotational 3D mechanogenomic Turing patterns of human colon Caco-2 cells during differentiation

**DOI:** 10.1101/272096

**Authors:** Gen Zheng, Alexandr A. Kalinin, Ivo D. Dinov, Walter Meixner, Shengtao Zhu, John Wiley

## Abstract

Recent reports suggest that actomyosin meshwork act in a mechanobiological manner alter cell/nucleus/tissue morphology, including human colon epithelial Caco-2 cancer cells that form polarized 2D epithelium or 3D sphere/tube when placed in different culture conditions. We observed the rotational motion of the nucleus in Caco-2 cells in vitro that appears to be driven by actomyosin network prior to the formation of a differentiated confluent epithelium. Caco-2 cell monolayer preparations demonstrated 2D patterns consistent with Allan Turing’s “gene morphogen” hypothesis based on live cell imaging analysis of apical tight junctions indicating the actomyosin meshwork. Caco-2 cells in 3D culture are frequently used as a model to study 3D epithelial morphogenesis involving symmetric and asymmetric cell divisions. Differentiation of Caco-2 cells in vitro demonstrated similarity to intestinal enterocyte differentiation along the human colon crypt axis. We observed rotational 3D patterns consistent with gene morphogens during Caco-2 cell differentiation. Single- to multi-cell ring/torus-shaped genomes were observed that were similar to complex fractal Turing patterns extending from a rotating torus centre in a spiral pattern consistent with gene morphogen motif. Rotational features of the epithelial cells may contribute to well-described differentiation from stem cells to the luminal colon epithelium along the crypt axis. This dataset may be useful to study the role of mechanobiological processes and the underlying molecular mechanisms as determinants of cellular and tissue architecture in space and time, which is the focal point of the 4D nucleome initiative.

## Background & Summary

We recently developed an automated 3D cell morphological classification allowing quantitative morphometry analytics of multiple geometric features, which are not accessible by traditional analysis of counting and signal intensity quantification [1]. Our shape analysis of nucleus and nucleoli of proliferating/non-proliferating fibroblast cells and epithelial/mesenchymal human prostate cancer (PC3) cells found morphology features including eccentricity and surface area to be useful predictors for these cell types. Symmetric and asymmetric cell divisions are the topological basis for the transformation from a single cell (a fertilized egg/ a stem cell) to a multi-cell system (an embryo/a piece of tissue). Our morphological classification protocol can potentially be applied to discriminating non-dividing cells/dividing cells as well as symmetrically dividing cells/asymmetrically dividing cells. Machine learning classifiers in our protocol can use topological characteristics describing patterned cells/nucleus and are directly applicable to quantifying the 4D nucleome in intact tissues. Analysis of actomyosin meshwork and cell shapes during embryo gastrulation which initiates asymmetric cell division and epithelial-mesenchymal transition also used eccentricity and surface area as a “small number of parameters” to describe the complex behaviour of the embryo tissue [2]. To help generate “proof of concept” data, in this study we chose a commonly used human colon epithelial cell model, Caco-2 BBe, tracked the formation of a coordinated epithelial cell sheet during differentiation of increased cells on a limited smooth, flat and hard glass surface. These results demonstrate the ability to model shape challenges of the 4D nucleome using morphometric analysis, which may be applied to coordinating multicell systems [3, 4]. Computational modeller with the goal to generate theoretical morphogenesis frameworks integrating the knowledge of mechanical, cellular and gene-regulatory levels may benefit from our previous work and this dataset with application suggestions [5–8]. Potential clinical applications in functional bowel disorders (FBS) including irritable bowel syndrome (IBS) and other colon diseases are also discussed [9–12].

## Data notes

Mechanobiology examines the role of physical and mechanical forces in the control of cell development and disease, in addition to chemicals and genes [13,14]. A major goal of the 4D nucleome project is to understand the subcellular details of this process compared to physical systems governed by Newton’s laws of motion [15]. The initial mathematical analysis of the multi-cell system using the digital computer based on Newton’s laws of motion proposed 4 types of changes involving the mechanical and the chemical parts:

(i) “The changes of position and velocity as given by Newton’s laws of motion.
(ii) The stresses as given by the elasticities and motions, also taking into account the osmotic pressures as given from the chemical data.
(iii) The chemical reactions.
(iv) The diffusion of the chemical substances. The region in which this diffusion is possible is given from the mechanical data.”

These processes are articulated in Alan Turing’s 1952 paper “The Chemical Basis of Morphogenesis” [16]. Based on these assumptions, Turing predicted specific shapes would be observed in biological systems at the cellular and tissue levels, e.g., “ring of cells” in 2D & gastrulation in 3D [2, 8, 17].

Rotational nucleus motion driven by actomyosin is observed within single cells [18]. Epithelial cells demonstrated rotational motion capabilities in 2 cell model, which was correlated with polarities [19] (Fig. 1-3). The rotational motion during 3D morphogenesis from a single cell to a 3D epithelium sphere were also detected [20]. The recordable sphere surface actomyosin meshwork curves looks like rubber bands demonstrating different eccentricity values (circle/ellipse/parabola/hyperbola) which are assumed to be generated by rotational motions might be used as basic elements for nucleus and cell morphology analysis (Fig. 4-6) [1, 2, 19]. Coordinated cells behavior and shape characterization emerged from 1 to 2 and multiple cells. However, new higher-resolution 3D/4D imaging and post-imaging analysis may be required to precisely track the coordinated 4D dynamics of patterned cells and nuclei.

Mechanical forces generated by the actomyosin network can force the cells and nuclei into different shapes [21, 22]. The forces from actomyosin network can be transduced to the genome via LINC (linker of nucleoskeleton and cytoskeleton) complex to the genome and induce epigenetic, and transcriptional changes [14, 23–25]. Human colon epithelial Caco-2 cells during the early stage of differentiation (2-4 days) demonstrated quantifiable changes of nuclear and nucleolar shape morphology detectable by our shape classification protocol [26]. Epigenetic modification H3K9me3 is sensitive to the mechanical forces, robust upregulation of H3K9me3 by mechanical force was observed within 1 min, and culture matrix stiffness can mediate H3K9 methylation [27, 28]. We observed the signature rotational pattern of H3K9me3 signals within the Caco-2 nucleuses coordinating with each other, where the signals seemed to be correlated with mechanical pressure spots (Fig. 7). Our shape classification protocol analyses the morphometry of nuclei and nucleoli that can be used as quantitative signature vectors for analysing the epigenetic modification shape variations between the cells, which are difficult to interpret using traditional methods based on homogenizing the cells (e.g. ChIP-seq) [1].

**Fig. 1.**
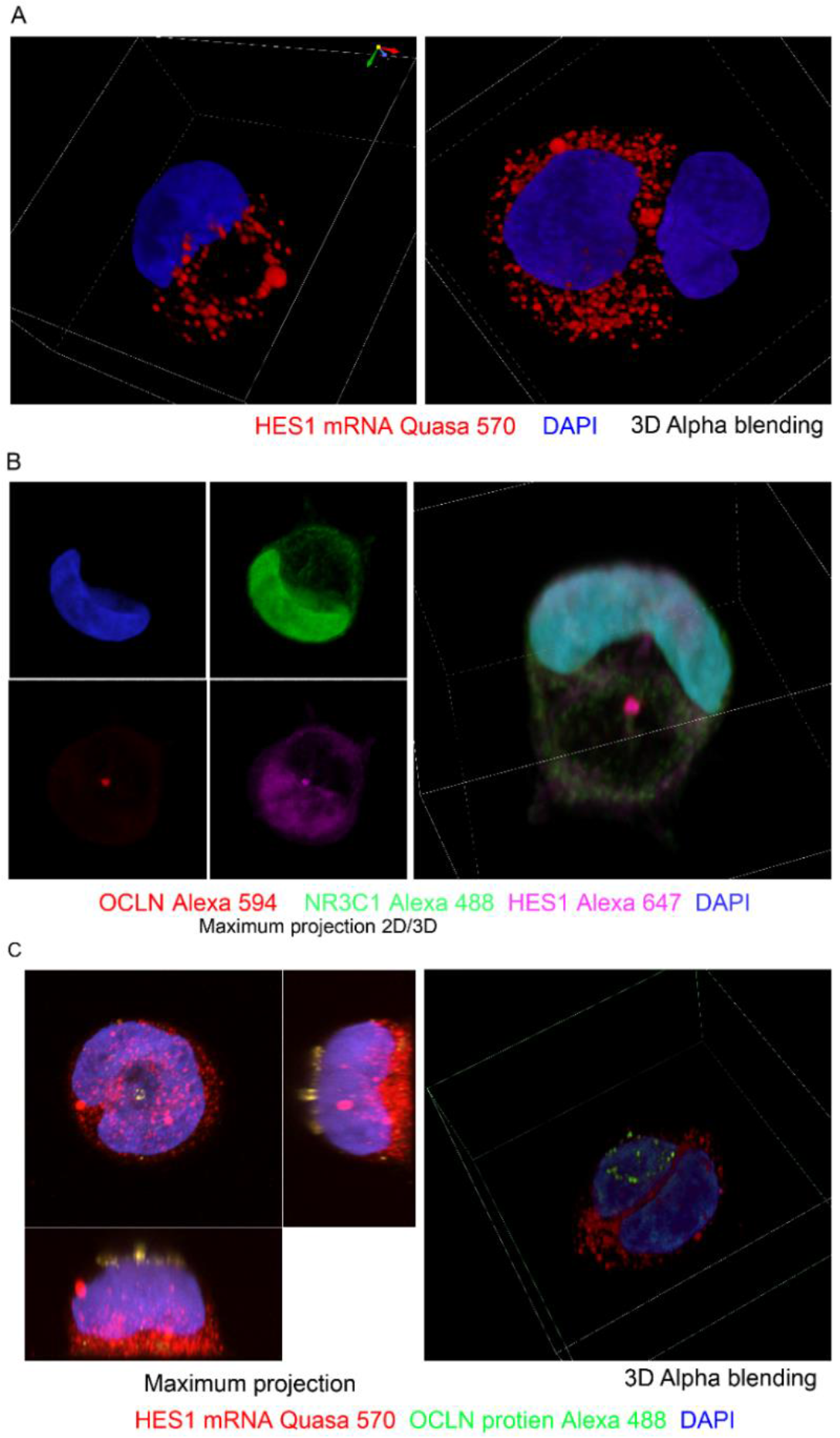
Ring patterns in 1-2 Caco-2 BBe cells at day 2. A. HES1 mRNA showed circular distribution in 1 cell and neighboring cells with or without HES1 mRNA are observed. B. Caco-2 BBe cells on day 1 cover clip are labelled with OCLN, NR3C1 and HES1 antibodies. Single cell genome with OCLN HES1 morphogen center. C. Caco-2 BBe cells on day 1 cover clip are labelled with HES1 mRNA Qausa 570 oligo FISH probes and OCLN antibody. Single cell genome labelled with DAPI can be found in a torus shape. OCLN protein showed circular wave pattern, this pattern may correlate with actomyosin force [36]. Formation of “ring of cells” proposed by Turing may start from 1 to 2 cells, the HES1 distribution may conform to “The morphogen pattern in a ring of cells as deduced by Turing” [17].

**Fig. 2.**
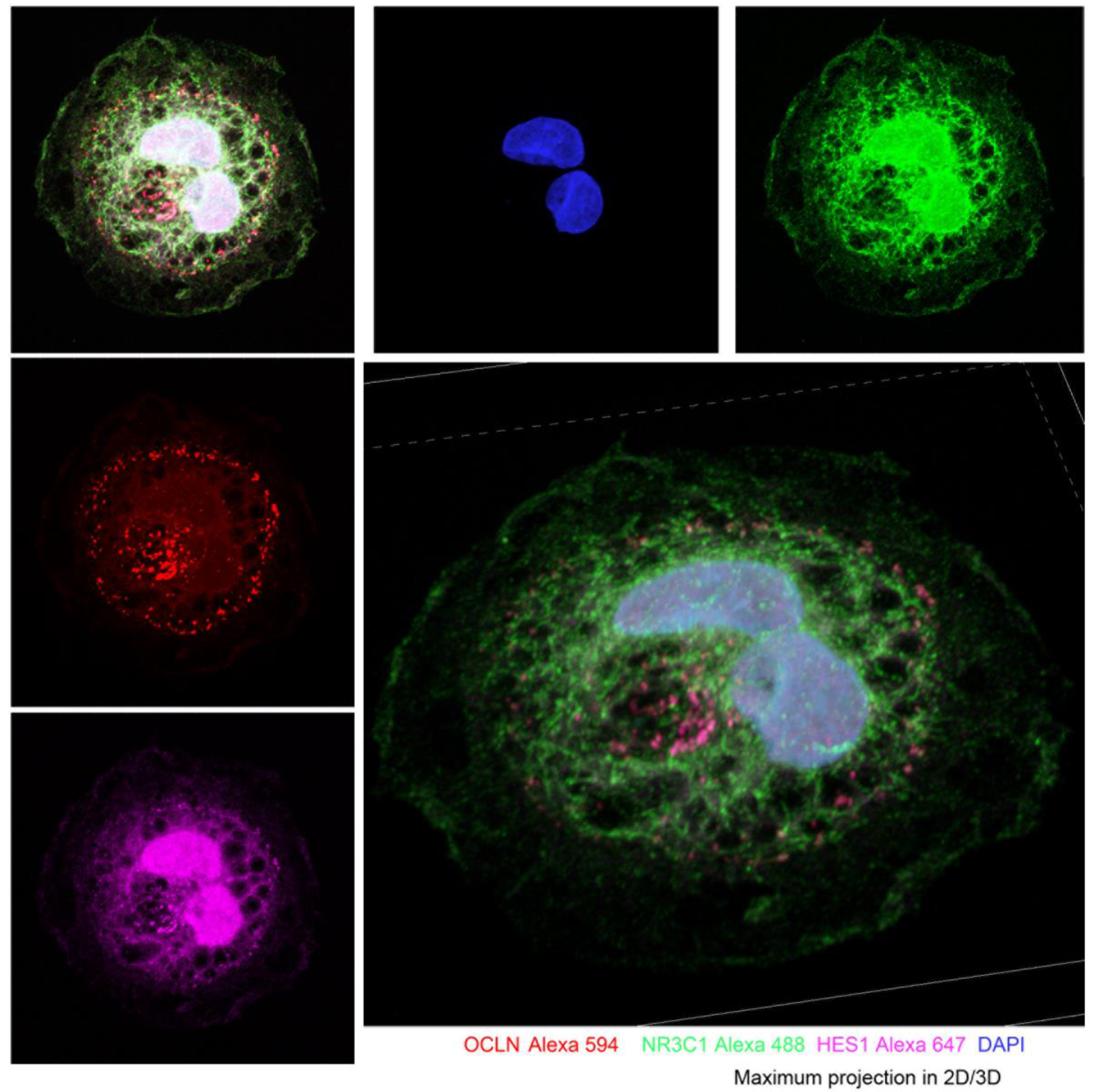
2 cells spreading spirally on flat 2D flat surface. 2 Caco-2 BBe cells on day 3 cover clip labelled with OCLN, NR3C1, and HES1 antibodies. Spirally spreading OLCN and HES1 can be seen. NR3C1 showed wave peak pattern comes from the spiral axis. OCLN’s circular wave pattern may correlate with actomyosin force [36]. The shape of the nucleuses looks conform to actomyosin Rho GTPase shape in a model of the rotational motion of polarized mammalian epithelial cells on micropatterns [19].

**Fig. 3.**
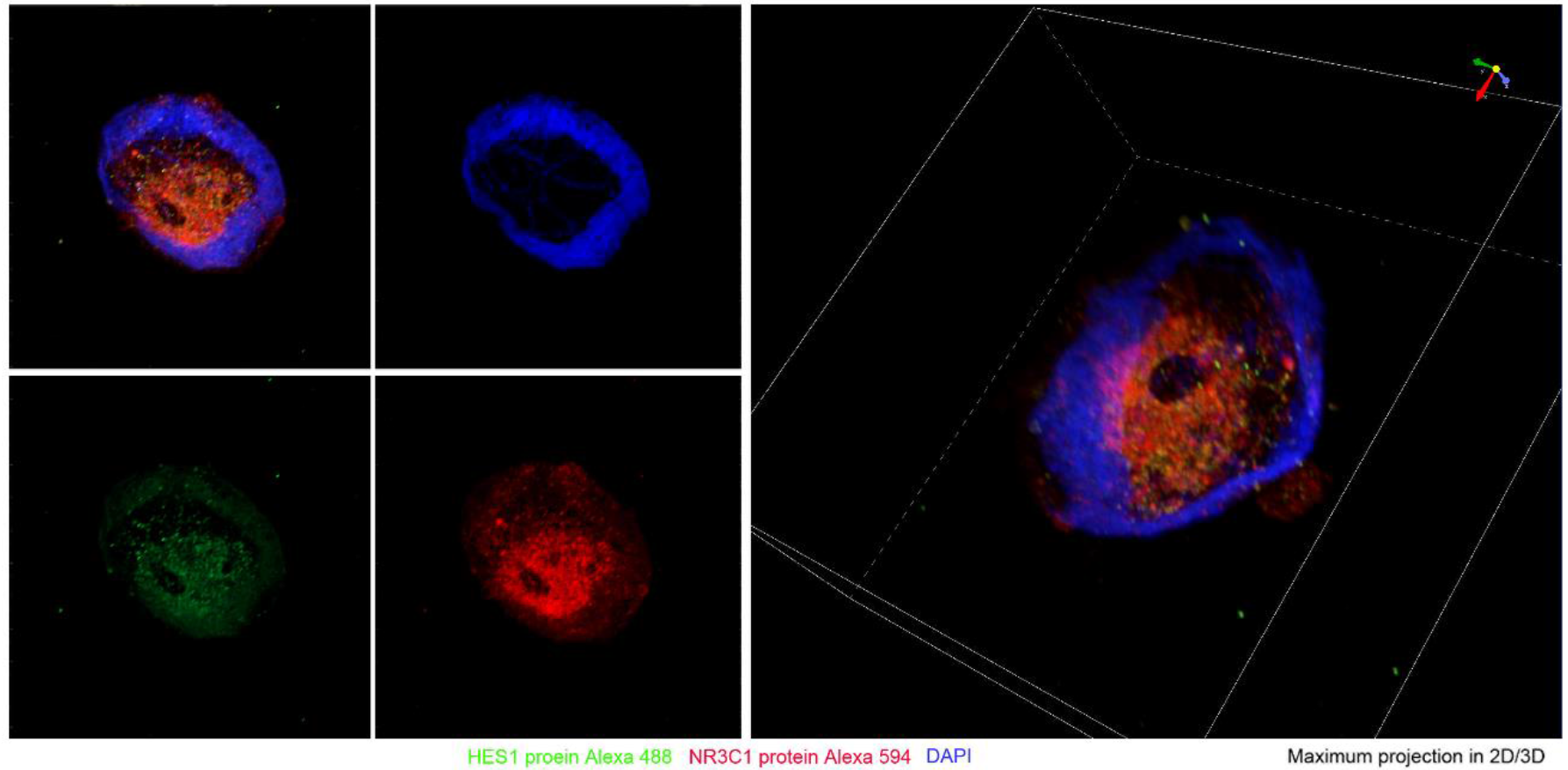
Vortex/Horn shaped cell with ring shaped genome. Caco-2 BBe cells on day 1 cover clip are labelled with NR3C1 and HES1 antibodies. DNA staining showed circular loop pattern, a hole can be seen located at the center of a vortex. Diffusion of the transcription factor NR3C1 and HES1 from the hole can be seen. This pattern may correlate with actomyosin force [36]. The shape of genome may conform to a “Hopf bifurcation” used to illustrate the mathematics of the genome [6].

**Fig. 4.**
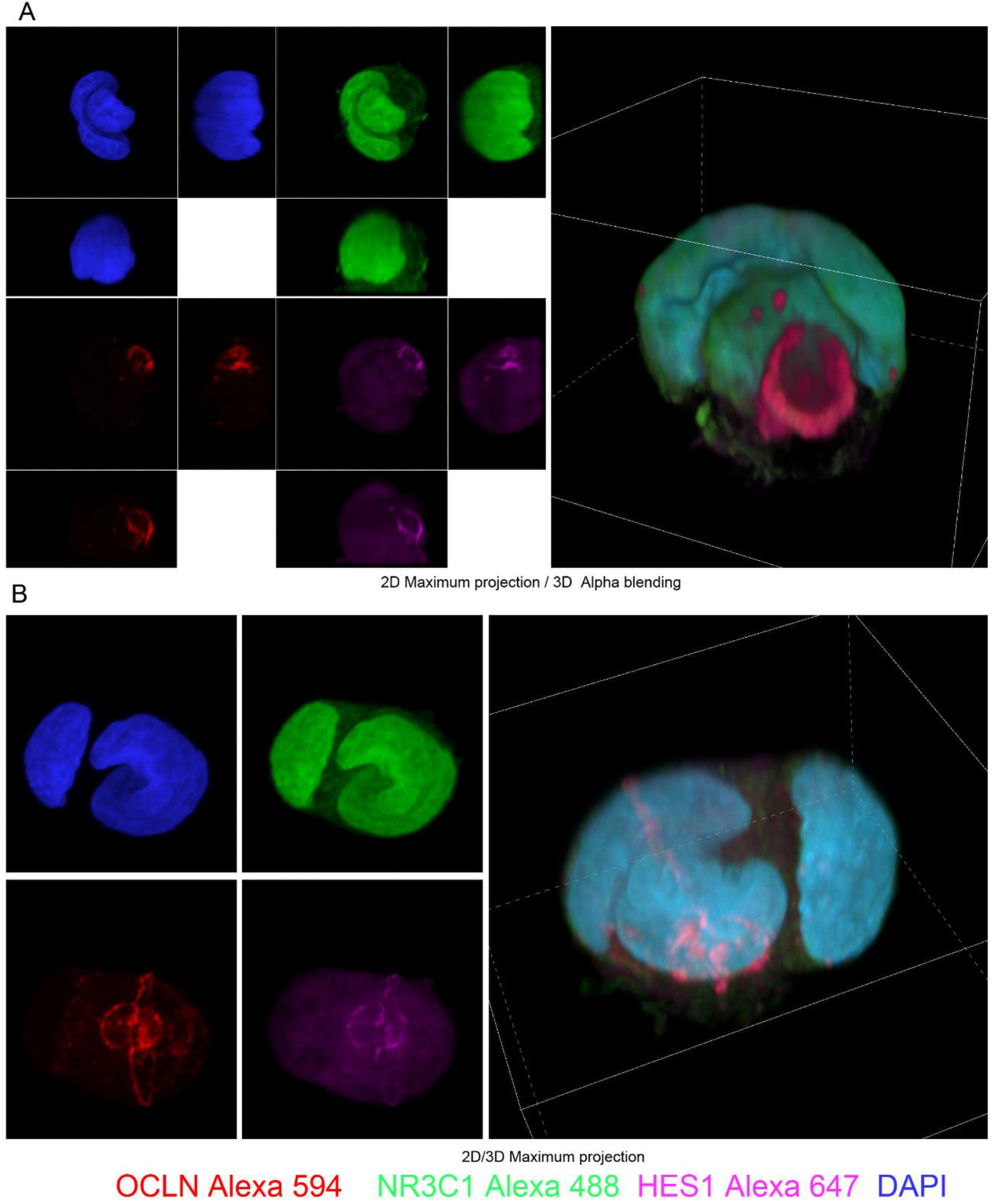
OCLN rings on sphere surface of the cells. Caco-2 BBe cells on day 1 cover clips are labelled with OCLN, NR3C1, and HES1 antibodies. OCLN showed ring patterns indicating different eccentricity values (circle/parabola/hyperbola) on the sphere surface of non-separated nucleuses.

**Fig. 5.**
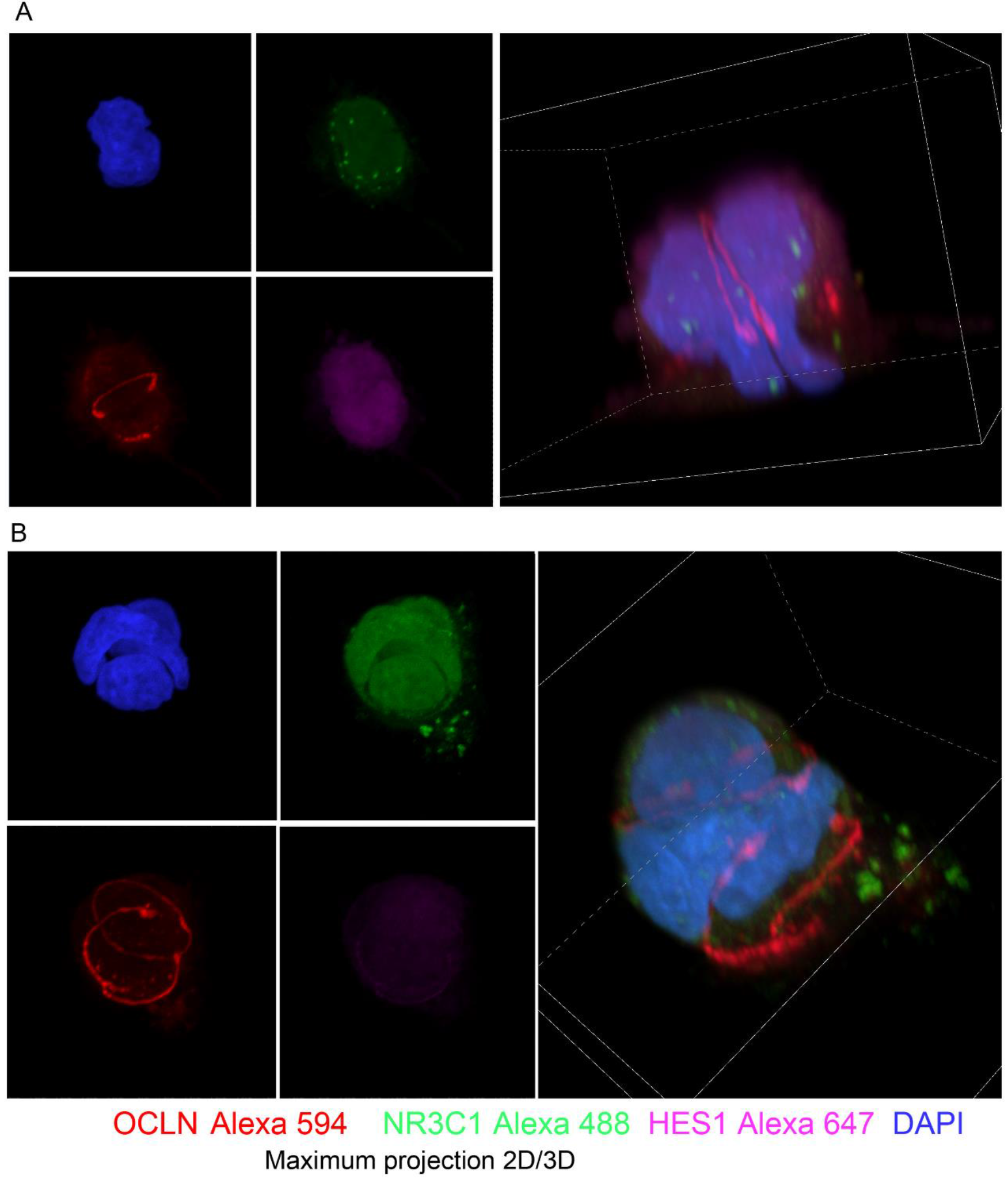
OCLN orbit tracks between separating nucleuses. Caco-2 BBe cells on day 1 (A) / day 3 (B) cover clips are labelled with OCLN, NR3C1, and HES1 antibodies. OCLN track showed bright ellipse orbits like pattern.

**Fig. 6.**
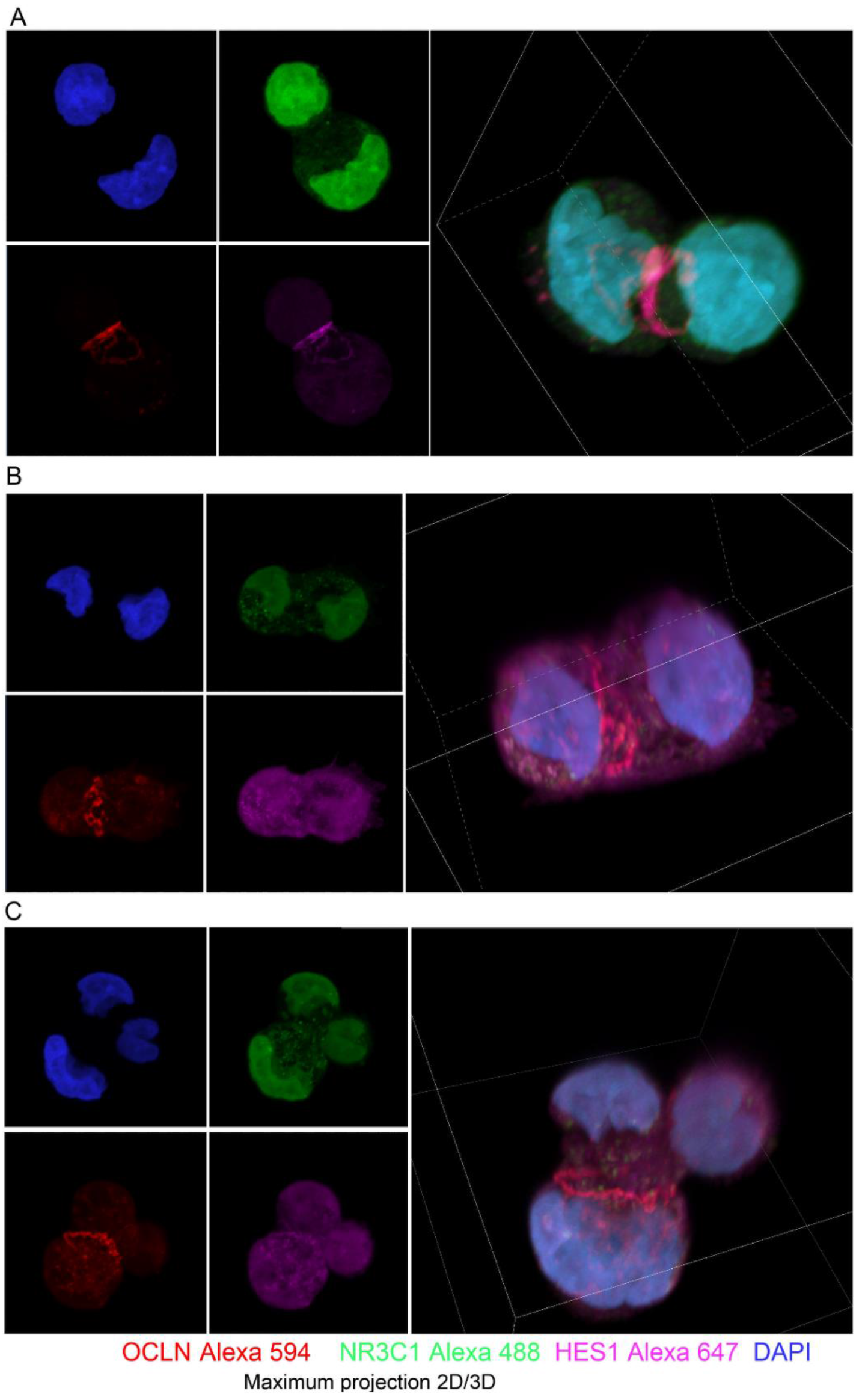
OCLN contractile ring between separated nucleuses. Caco-2 BBe cells on day 1 cover clips are labelled with OCLN, NR3C1, and HES1 antibodies. OCLN showed contractile ring pattern separating the cells with separated nucleuses. The shape of the nucleuses looks conform to actomyosin Rho GTPase shape in a model of the rotational motion of polarized mammalian epithelial cells on micropatterns [19].

**Fig. 7.**
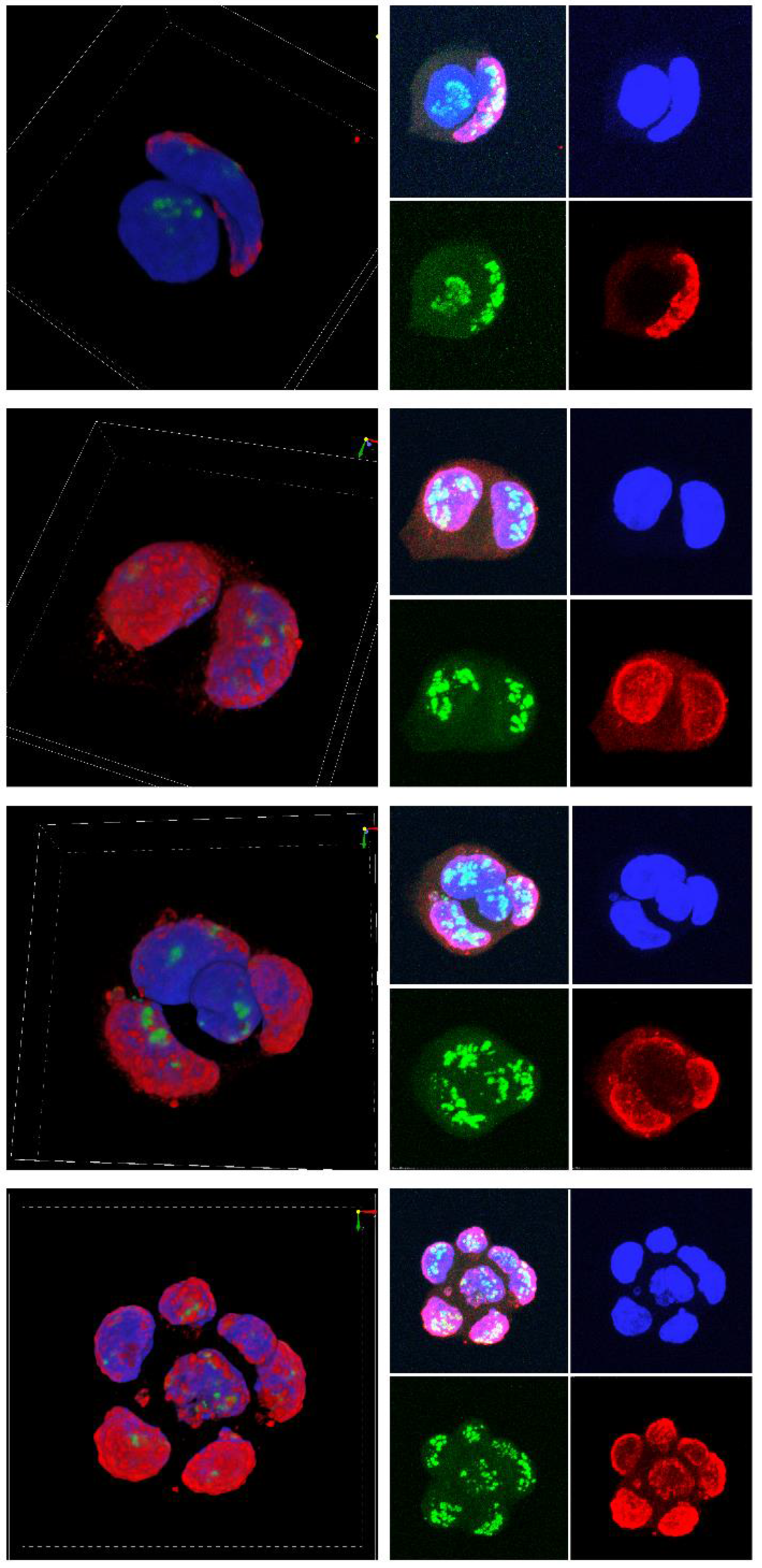
H3K9me3 & nucleoli in nucleuses. Caco-2 BBe cells on day 1 / day3 cover clips are labelled with H3K9me3 and fibrillin antibodies. Rotational patterns of H3K9me3 can be seen, there might be a correlation of H3K9me3 with mechanic forces [27].

Mechanical crosstalk between cells via apical tight junctions of epithelial cells membrane mediate the functional signal transduction and pathogenesis [29–31]. Caco-2 is a frequently used cell model for epithelial mechanobiology/morphogenesis. It can form: (1) 2D epithelial membrane; (2) 3D epithelial sphere/cyst/tube; or (3) dynamic membrane with villin structures close to *in vivo* gut epithelium with different mechanobiology culture conditions [20, 32–34]. The 2D culture of this model has been used for decades to study gut barrier function and paracellular permeability dependent on the expression and function of epithelial cell tight junctions [35]. These were also mathematically validated as a mechanobiology model of 2D Turing patterning. Self-organizing actomyosin meshwork was observed and simulated to drive the oscillatory patterning the cell cortex at epithelial apical tight junctions, the actomyosin forces demonstrated circular wave pattern [36]. Actomyosin oscillation is important in tissue-level cell-cell coordination including the epithelium [37]. The 3D culture of a single Caco-2 cell to a polarized epithelial sphere is also frequently used to study the molecular mechanism of epithelial morphogenesis involving symmetric and asymmetric divisions via various interventions [20, 38–40]. The actomyosin network is responsible for the morphology of the 3D sphere/cyst formation of Caco-2 cells [32]. Rotational motion during the 3D morphogenesis may correlate with the formation of the Turing-proposed “ring of cells”/dome observed on flat glass coverclips (Fig. 8-9) [16, 20]. The formation of “rings of cells”/domes are also observed in functional enterocyte epithelium differentiated from human induced pluripotent stem (iPS) cells, which could be used to replace the Caco-2 cell model. Wound healing morphogen FGF2 treatment, the latter inhibits differentiation and prevents the formation of “ring of cells”/dome in Caco-2 cells [8, 41–43]. Knockdown extracellular matrix protein which stimulates differentiation in Caco-2 cells delayed the “ring of cells”/dome formation [44]. Apical tight junctions are also involved in maintaining the polarity and actomyosin network controlled “dividing angle” of the epithelial cells in 3D morphogenesis [45]. Tight junction containing cellular structure midbodies are recognized as the organelle regulating symmetric division and asymmetric division which determine the cell fate [46] (Fig. 6). Rotational motion-generated “Yin-yang (YY)” shape has been proposed to correlate with the mammalian cell symmetry breaking which is “essential for cell movement, polarity, and developmental patterning” [19, 47]. We observed large fractal YY shaped cells in our study (Fig.10 and Supplementary Movie.S1).

**Fig. 8.**
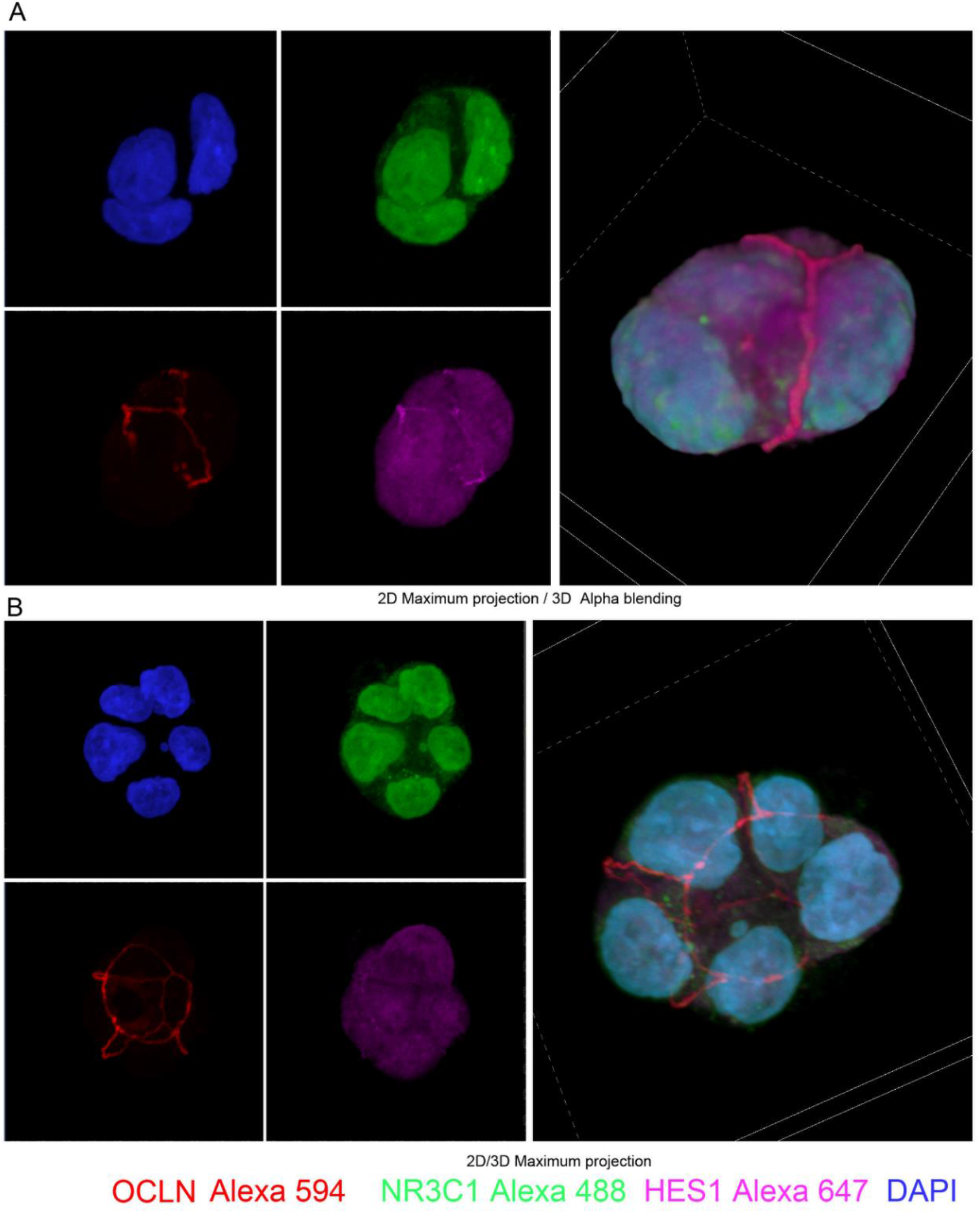
Tendency of forming 3D structures. Caco-2 BBe cells on day 1 cover clips are labelled with OCLN, NR3C1, and HES1 antibodies. Cells showed 3D organization, OCLN showed 3D orbit tracks like pattern. This pattern may correlate with rotational motion observed during 3D morphogenesis of epithelial cell morphogenesis [20]. Euler characteristic should be considered.

**Fig. 9.**
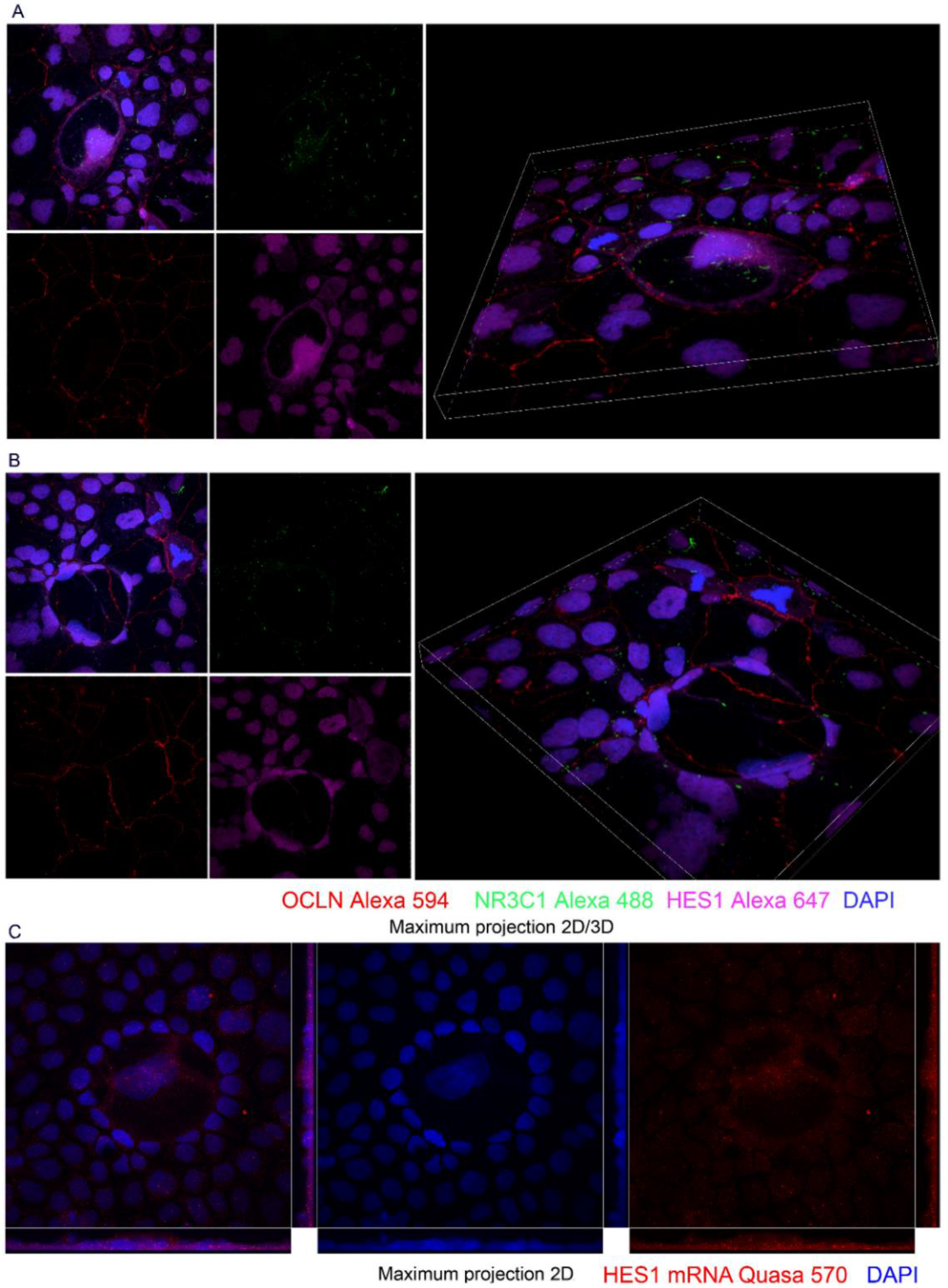
Circular spreading cells on flat 2D surface. Caco-2 BBe cells on day 4 cover clips are labelled with OCLN, NR3C1, and HES1 antibodies. Formation of “ring of cells” may conform to “The morphogen pattern in a ring of cells as deduced by Turing” when HES1 is considered as the morphogen [17]. A. Spirally spreading HES1 wave toward OCLN ring from the nucleus. B. HES1 protein ring guided ring of cells. C. HES1 mRNA ring guided ring of cells, HES1 mRNA showed bridge like pattern within the ring.

**Fig. 10.**
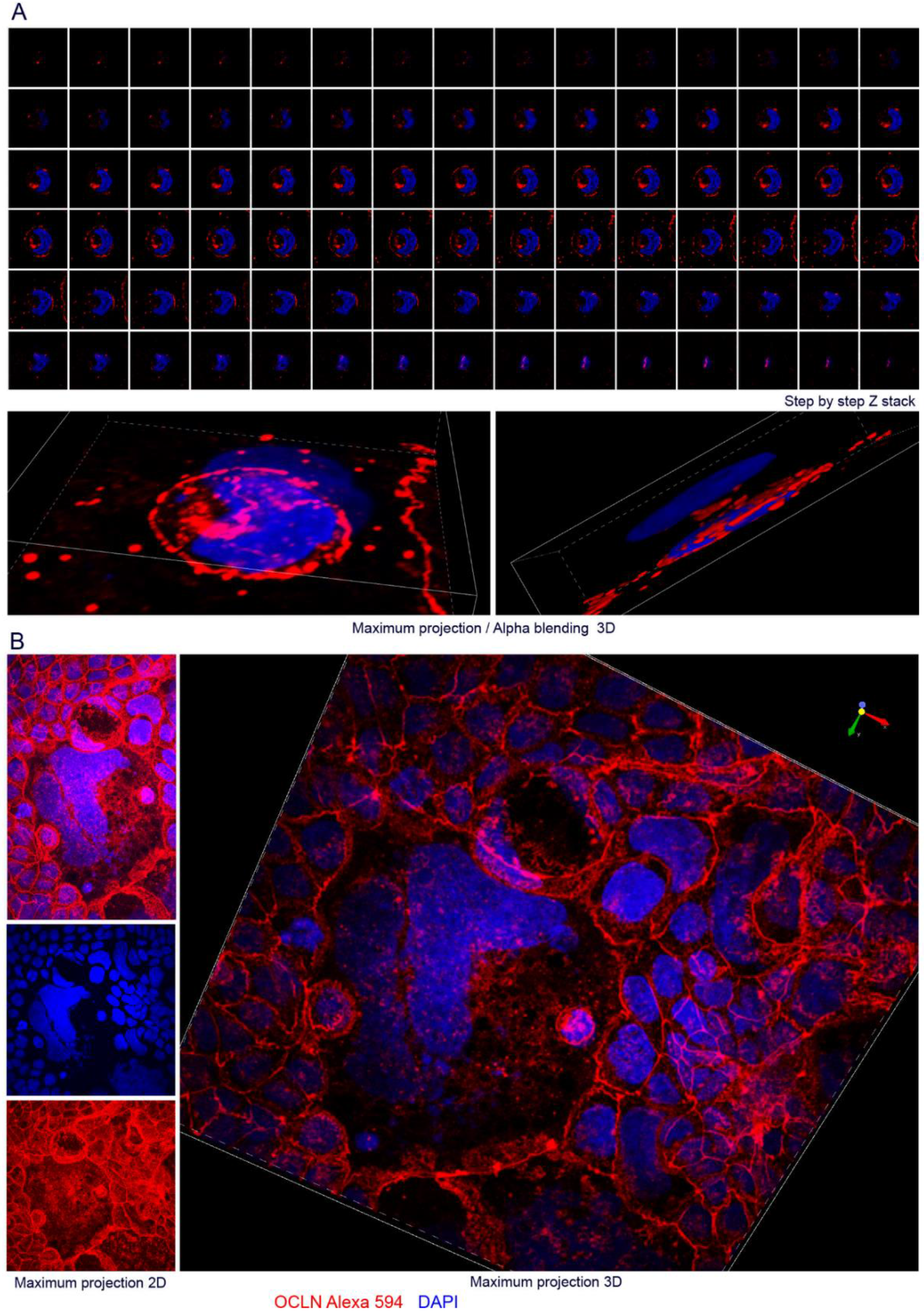
Yin-yang (YY) shapes, small YY in big YY. Caco-2 BBe cells on day 6 cover clips are labelled with OCLN protein and DNA. YY shape may correlate with rotational motion and symmetry breaking [19, 47].

Mechanical forces can trigger biochemical Notch signaling pathway which was considered sensor of mechanobiology signals. These mechanical forces pull cells apart leading to proteolytic activation of Notch while tight junctions holding cells together apparently counteract against the pull [14, 48, 49]. This process contributes to Notch signaling pathway’s ability to integrate multiple micro-environmental signals and sense cell-cell interaction, trigging the downstream Notch signaling target HES1 transcription [50, 51]. Thus, cell shapes can affect the Notch signaling triggered events [52]. Key transcription factors/”master” regulators are considered important 4D nucelome modulators [5, 8, 15]. Glucocorticoid receptor (NR3C1) and transcription factor HES1 can bind their own promoters for dynamic autoregulation, they can also repress each other’s transcription via binding to the promoters [53, 54]. This special “hard-wiring” matches with “a two-gene network with two repressors” model supports Turing’s patterns [6, 7]. We found that these motifs displayed opposite distribution along Caco-2 cell differentiation with increased cell density and human colon crypt axis, which could be simulated by calculating forces between the cells via cell shapes [5, 55]. This pattern conforms to “a stable equilibrium as a cell type” though differentiation in a mathematical model of the genome which include the toggle gene circuits (from high averaged surface area HES1+/NR3C1-PRC2&H3K27me3+ to low averaged surface area HES1-/NR3C1+ PRC2&H3K27me3−) [6, 56]. HES1 is an extensively studied and mathematically simulated oscillatory transcription factor, it is also involved in 2D epithelium Turing patterning [57, 58] (Fig. 1,11). Recent mathematical models confirmed that HES1 promoter is the crosstalk hub in Notch-Wnt interactions of the intestinal crypts, our recent study also suggested that physiological glucocorticoid (GC) signal can affect HES1 expression via HES1 promoter negative glucocorticoid response element (GRE) [5, 59, 60]. We observed down-regulation of HES1 by psychological stress level of glucocorticoids, HES1 deficiency resulted increased goblet cell population, altered colon crypts morphology and impaired gut homeostasis which conform to the mathematical model [5, 9, 59]. These features indicate that the HES1 promoter is also a potential crosstalk hub of Notch-GC interactions in colon crypts [5, 60]. The levels of these opposite transcription factors are coupled with tight junction protein levels. We observed robust down-regulation of gene oscillator HES1 and robust up-regulation of tight junction protein CLDN1 correlated with NR3C1 upregulation during the Caco-2 cell differentiation. These features may correlate with the morphogenesis of intestinal crypts structure [8]. There might be a hardwired antagonism between tight junction CLDN1 gene and HES1 gene coded close on chromosome 3 in opposite directions too. Their promoter positive/negative GREs and HES1 binding N-boxes provided regulatory elements required for dynamic regulation [5]. “Hardwiring” of HES1-CLDN1 & NR3C1 may meet the proposed criteria of the repressilator in a mathematics model of the genome [6, 7]. Notch’s mechanical feature and these transcription factor/promoter hardwiring provided potential hardwired linkage between oscillatory actomyosin force and oscillatory gene transcription in the scenario of morphogenesis [49, 51, 61, 62]. We hypothesize that HES1-CLDN1 & NR3C1 genes could be candidates to test gene morphogen hypothesis and investigate the 4D nucleome dynamics during differentiation and morphogenesis [5, 6, 63]. The microtubule-dependent global mRNA polarization is recently verified in the intestinal epithelium. This is consistent with our observation of polarized HES1 mRNA distribution [64] (Fig.1 & 11). Asymmetric distribution of HES1 is also found paired with asymmetric H3K9me2 histone modification in the scenario of asymmetric cell division, they may be used the markers in identifying proliferating/differentiating and apoptosis cells along the colon crypt axis [65] (Fig. 1,7,11). These markers may help us further analyze the topology context of the patterning multi-cell system.

**Fig. 11.**
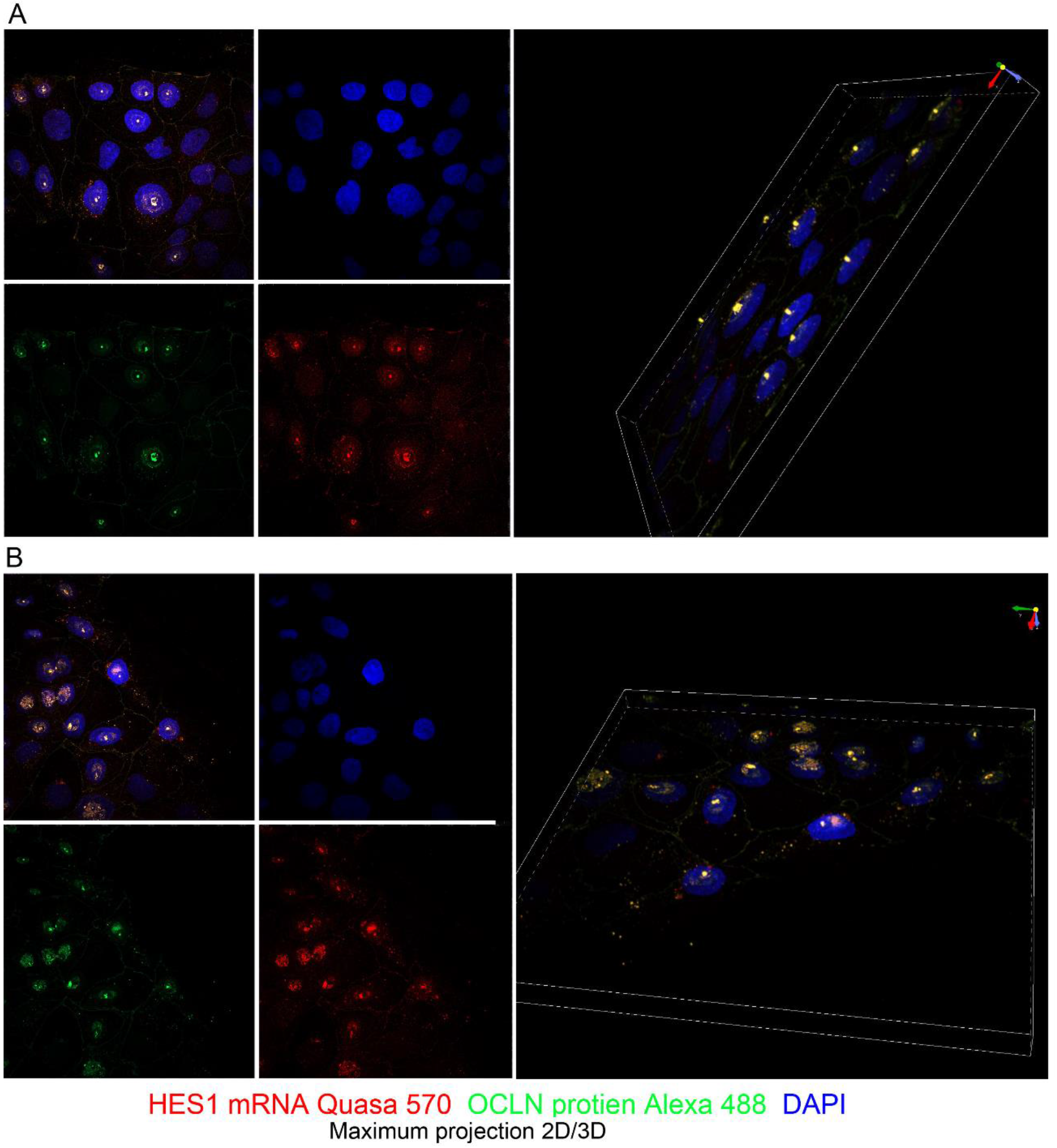
Turing pattern of cells with or without HES1 mRNA / OCLN protein spirally spreading from pole of cell polarization like circular waves. Caco-2 BBe cells on day 4 cover clips are labelled with HES1 mRNA and OCLN protein. HES1 on and off pattern is similar to what was observed in colon crypts which conform to Turing patterning [59]. Circular wave shaped HES1 mRNA and OCLN protein can be observed.

**Fig. 12.**
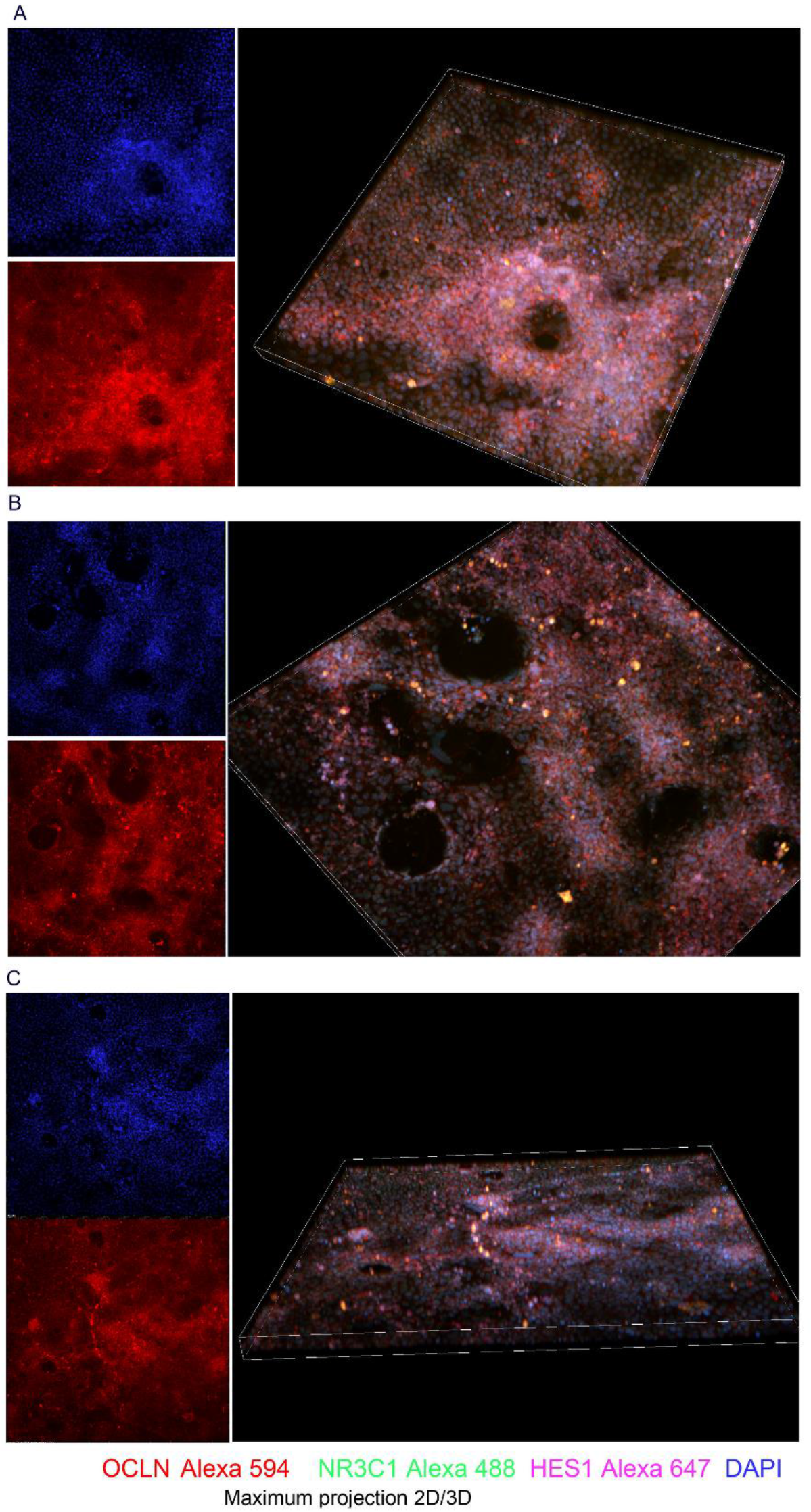
Huge rings of cells and circular waves of cells. A. Caco-2 BBe cells on day 21 were treated with vehicle for 2 h, cover clips are labelled with OCLN, NR3C1, and HES1 antibodies. B, C. Caco-2 BBe cells on day 20 were treated with 500 nM GC for 24 h, cover clips are labelled with OCLN, NR3C1, and HES1 antibodies.

Crosstalk between the brain-gut axis and microbiome is intensely involved in functional bowel disorders (FBDs) including irritable bowel syndrome (IBS), latest reports suggest that the key transcription factor HES1 & NR3C1 played pivotal roles in bacteria-host communication in the scenario of FBDs [5, 9–11]. The real-time 4D crosstalk between the gut microbiome and Caco-2 epithelium has been studied in a human gut-on-a-chip system able to apply mechanical forces [66–69]. Improvement of probiotics utilizing the 4D nucleome algorithms should be very promising in generating personalized probiotics against FBDs [1, 6, 10, 11, 15, 70–73].

Actomyosin meshwork used to demonstrate morphogenesis can be tracked with CLDN1 binding peptide employing a preclinical confocal colonoscopy system suggesting the potential application correlating geometry information and pathological colon lumen tissue patterns [8, 12, 74, 75]. Human colon epithelium demonstrates various patterns that correlate with different physiological and pathological conditions under colonoscopy, which can be potentially identified using recently developed artificial intelligence algorithms, the biological mechanisms underlying these patterns remain unknown [12, 76]. 2D/3D patterns consistent with Turing’s hypothesis appear to be present in our clinical data (Fig. 13). The colon lumen pattern looks similar to self-organizing spots under the skin, morphogenetic feedback mechanisms are suggested to regulate the topology architecture of tissues [8, 77]. One may speculate that application of mechanobiology culture methods to isolated human colon crypts maintained in a matrix environment will replace the 2D Caco-2 cell model [78]. We anticipate that application of the programmable “gut-on-a-chip” mechanobiology culture methodology to 2D cultures will demonstrate epithelium morphology textures similar to the human colon lumen surface with functional microbiome, this system can be used to study the topology of colon epithelium in response to morphogens including FGF2/glucocorticoids/microbiome etc. which are currently used clinically [10, 66, 79–81]. Recent clinical studies demonstrated that *Helicobacter pylori* infection and eradication can induce endoscopic morphological changes and epigenetic changes in samples collected, these trends may also exist in colon [75, 82, 83]. Patient colon crypts can be isolated from the biopsies for actomyosin meshwork/4D nucelome analysis and organoids culture (Fig. 14) [8, 63, 84]. Study of movement of specific genes within the nucleus has been proposed and recently performed with 4D nucelome methods, human colon crypts could be used as the film recording the life from stem cells to differentiated cells [63, 85]. Correlating morphogenesis with oscillatory gene expression, oscillatory gene movement, super-resolution periodic patterns of chromatin and structural perspective of cooperative transcription factor binding (HES1 & NR3C1 for example) which biophysically shape DNA double helix in poloidal and toroidal directions will be required to generate convincing proof of gene morphogen hypothesis following Alan Turing’s “NON-LINEAR THEORY. USE OF DIGITAL COMPUTERS” legacy guidance [8, 16, 63, 86–88]. Deliberate mathematical planning is required to efficiently utilize the electronic health record (EHR), endoscopy observation/procedure, functional magnetic resonance imaging (fMRI),-Omics analysis, organoids *in vitro* culture and 4D nucelome datasets to guide the next generation of research of FBDs targeting the “Gut Microbiota-Brain Axis” [8, 11, 70, 89–92].

**Fig. 13.**
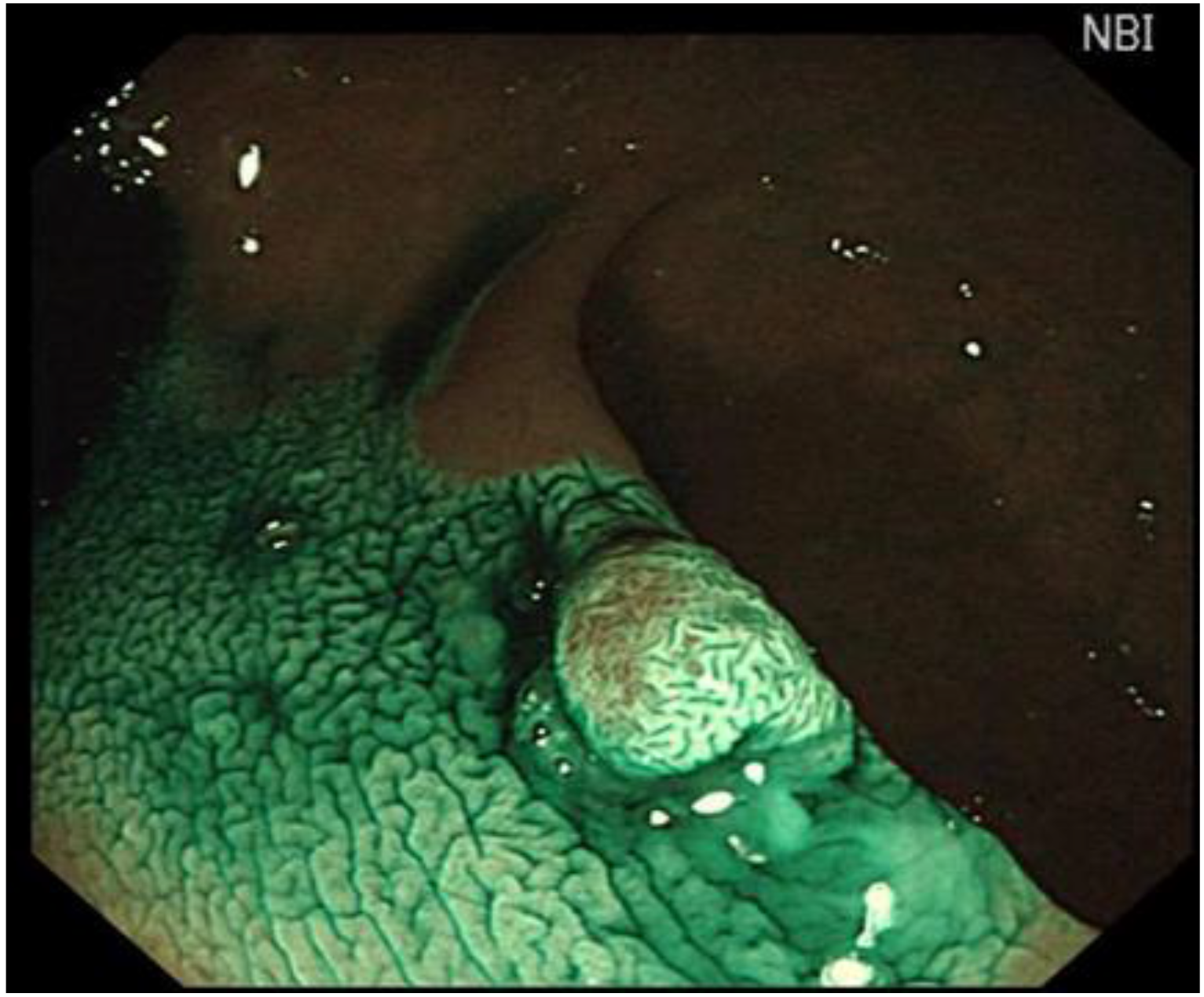
An example of colonoscopy image containing 2D/3D Turing patterns. The texture of colon surface showed similarity to 2D/3D Turing patterns [95, 96].

**Fig. 14.**
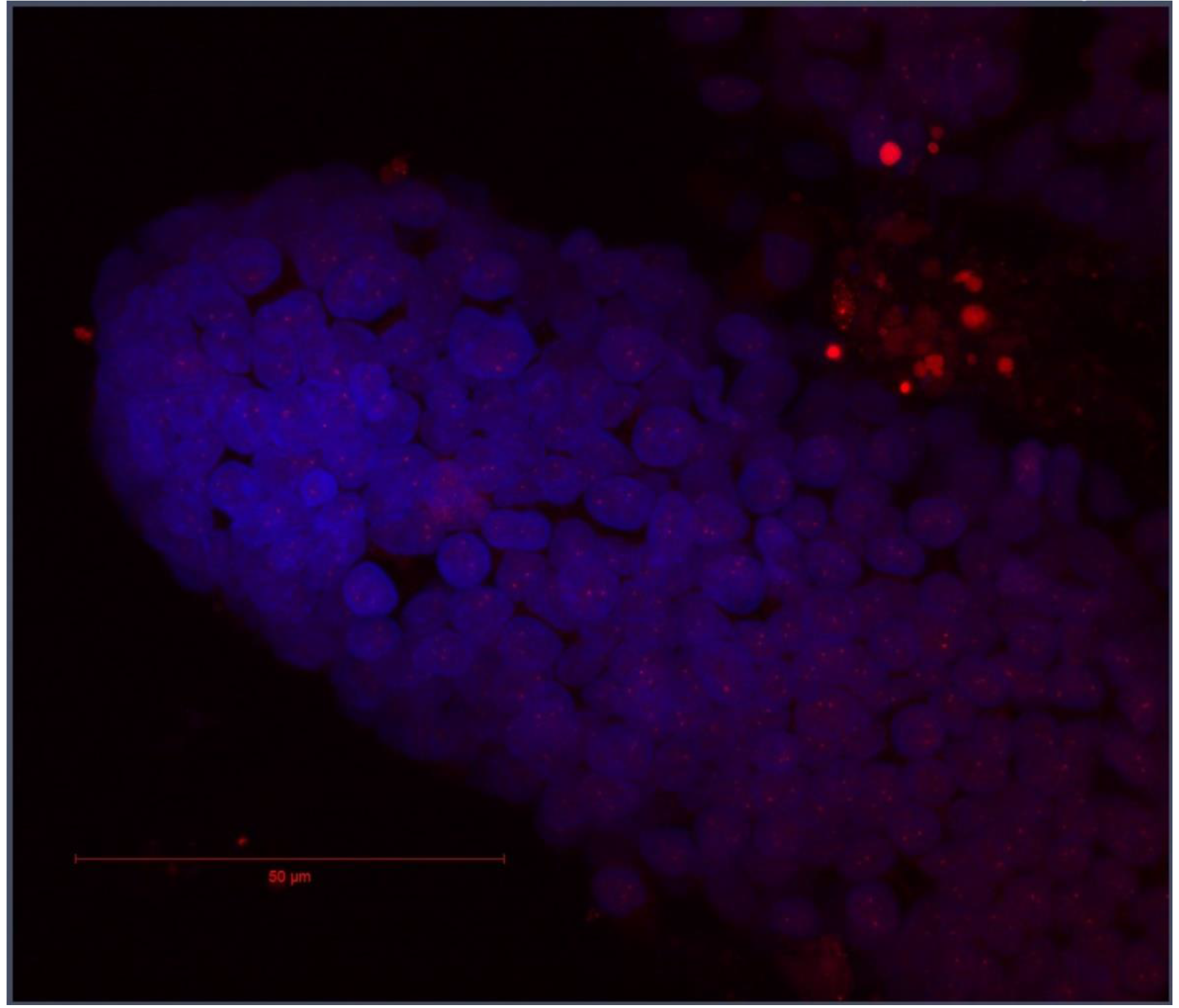
An example of BAC-FISH analysis of 3D human colon crypts isolated from clinical biopsy.

The use of data sets similar to what has present in this communication combined with evolving advanced analytical techniques will enhance our ability to derive novel information regarding the determinants of cell morphology and tissue patterns in physiologic and pathophysiologic conditions. The computational modeller may want to consider nucleuses as “cellular decision-making” units in “cellular decision-making networks” with gut intelligence because colonic crypts/Caco2 cells are validated biomedical engineering models [93]. Computational modelling cellular decision in response to multiple biochemical and biophysical cues may need mathematics able to handle multiple dimensions [93]. The advanced mathematical technique of shaping differential topology, dynamical systems, bifurcation, and chaos etc. will be necessary in addition to “the relatively elementary mathematics” used by Alan Turing [6, 7, 16]. “Non-linear equations will be used.” might be considered as “Turingology bombe” conjecture proposed for cracking the Enigma of morphogenesis by the legendary cryptologist [16].

## Methods

### Caco-2 BBe cell culture

Caco-2 BBe cells are kindly provided by Prof. David E. Smith, Department of Pharmaceutical Sciences, and the University of Michigan. Cells were cultured with DMEM suppled with 0.01 mg/ml human transferrin; 10 % FBS in humidified 37 °C incubator with 5 % CO_2_. Cells were seeded at 50-60 % density, the medium was replaced every 2 days. These methods are expanded versions of descriptions in our related work [5].

### Fixation

ZEISS 0.17 mm coverclips was cleaned with cleaning solution (50% ddH_2_O, 25% ammonia and 25% Isopropanol) with 5 min agitation and washed with ddH_2_O 5 times 3 min agitation. Then the autoclaved and dried coverclips were put into non-coated 8 well plates before seeding the Caco-2 BBe cells. Fixation followed these steps: 1. 1× PBS wash twice; 2. 4% PFA/1× PBS for 10 min at room temperature; 3. 0.5% Triton-X100 in 1× PBS for 15 min; 4. 20% glycerol in 1× PBS treatment for at least 30 min; 5. 3 times repeated freeze/thaw in liquid nitrogen followed by soaking in 20% glycerol/PBS with a carbon tipped reverse tweezer; 6. 1 time wash in 1× PBS; 7. Incubate in 0.1 M HCl for 5 min; 8. Rinse in 2× SSC 9. Incubate in 50% formamide/2×SSC (at least 30mins at RT). Samples in 50% formamide/2×SSC kept in dark at 4 °C could be used 1 month after fixation. For RNA FISH, RNase free ddH_2_O/ 10×PBS/ 20×SSC were used. These methods are expanded versions of descriptions in a related work [94].

### Immunofluorescence (IF)

Samples were washed once with 1× PBS and blocked with 2% BSA in PBS for 30 min at room temperature, labeling with blocking solution diluted antibodies was performed overnight at 4 °C, then the samples were washed 3 times with PBS and mounted with Prolong Gold with DAPI (Thermo Fisher). For optimized DNA staining, DAPI (Thermo Fisher) in PBS 10 min staining following manufacture’s manual was recommended especially when cells are dense, Prolong Gold without DAPI should be used for this option. HES1 Alexa 647 (1:200; Abcam), NR3C1 Alexa 488 (1:150; Cell signaling) and OCLN Alexa 594 (1:600; Thermo Fisher) are used for images shown.

### Oligo FISH of mRNA

Stellaris^TM^ predesigned human HES1 mRNA oligo FISH probe was bought from LGC Biosearch, samples are washed once with wash buffer (2× SSC, 10% formamide) and labeled with manufacturer’s protocol. Prolong Gold with DAPI could be used for easy handling and DAPI staining followed by Prolong Gold without DAPI mounting could be used for better DNA staining. OCLN antibody (EMD Millipore) and Alexa 488 secondary antibody (Thermo Fisher) are used in the pictures shown.

### 3D confocal microscope imaging

Nikon A-1 confocal microscope equipped with Diode based laser system for 405,488,561, and 640 excitations and operated with Nikon’s Elements software in Microscopy & Image Analysis Laboratory of University of Michigan Medical School was used for this study. 1K×1K images were taken with 0.1 μM Z-stack (for the 60× objective lens) or with 0.2 μM Z-stack (for the 10× objective lens) steps covering all the signals to maximize details from the Z axis. 3D data sets are processed with Nikon’s Elements software or the viewer on a 2K or 4K screen. Satisfying screenshots are clipped and shown.

### Data Records

Caco2-BBe cells are seeded on flat and smooth hard glass coverclip surface and fixed at serial time points. We chose the fixation method compatible with DNA(BAC-FISH), RNA (Oligo-FISH) and immunofluorescence labelling to track DNA/RNA/protein forms of genes. We chose a fixation method tested compatible for DNA(BAC-FISH)/RNA(Oligo-FISH)/Protein(IF) labelling and the samples can be stored for over 1 month for fixing the samples during the 21-day differentiation and labelled the Caco-2 cells with tight junction protein occludin and transcription factor HES1 and NR3C1. Cells are grown on the smooth hard glass surface and cannot form 3D spheres without 3D matrigel matrix support, we hypothesize that the actomyosin force driving the formation of 3D spheres generated the rotational patterns we have shown. We chose those interesting “geometrically informative” sight fields and recorded with confocal Z-stack images. This dataset may be useful to biophysicists, imaging data scientists, and mathematicians to test the applicability of “imaginary biological systems” to real biological systems following Alan Turing’s legacy guidance.

Imaging protocols, original and segmented data, and the source code are made publicly available on the project webpage: http://www.socr.umich.edu/projects/3d-cell-morphometry/data.html

## Acknowledgements

We acknowledge Microscopy & Image Analysis Laboratory (MIL), the University of Michigan Medical School for providing the core service. This study was supported by NIH RO1DK098205 (JWW), NIH R21AT009253 (JWW/SH), NSF grants 17348531 and 1636840, and NIH grants P20 NR015331 and P50 NS091856.

## Author contributions

G.Z. conceived the experiments shown in Fig. 1-12 Fig.14.

S.Z. acquired image shown in Fig. 13.

W.M. acquired image shown in Fig. 14.

A.K., I.D.,W.M., S.Z. & J.W. participated discussion and editing.

All authors reviewed the manuscript.

